# Individual-level social associations and social information use predict foraging success in winter mixed-species flocks

**DOI:** 10.64898/2026.10.23.761426

**Authors:** Faiza Hafeez, Laura N. Vander Meiden, Aidan E. Hand, Allison E. Johnson, Stella F. Uiterwaal, Sagan R. Smith, Daizaburo Shizuka

## Abstract

The acquisition of social information to improve foraging efficiency is considered one of the primary drivers of mixed-species flocks (MSFs). This flow of social information occurs among individuals of multiple species, but analyses of MSFs at the level of species fail to capture the nuances of this dynamic. We investigated how individual-level social associations in North American MSFs impact discovery of novel feeders presented alongside vocal cues. We used radio-frequency identification (RFID) data loggers to track arrivals of individually tagged black-capped chickadees (*Poecile atricapillus*), white-breasted nuthatches (*Sitta carolinensis*), and downy woodpeckers (*Dryobates pubescens*) at fixed feeders, and during novel feeder trials paired with chickadee, nuthatch, or white noise playbacks. For chickadees, a higher number of total associations and conspecific playbacks improved novel feeder discovery. The foraging benefits in nuthatches and woodpeckers were predicted by their individual-level associations with chickadees and these benefits only manifested during chickadee playbacks. We demonstrate that the advantages of mixed-species flocking are not uniformly distributed across species; rather, they are contingent upon individually maintained social associations and the presence of social information. These findings highlight the necessity of quantifying individual variation to accurately assess the evolutionary drivers and fitness consequences of species interactions.

**Significance statement:** While there is increasing evidence that organisms can gain benefits from associating with heterospecifics in many contexts, most studies fail to account for individual-level variation in the costs and benefits of such species interactions. Here, we demonstrate that quantifying fitness consequences of mixed-species social associations at the individual-level is essential to understanding complex interspecific relationships and mechanisms underlying community assemblages. Using a network approach to study mixed-species flocks of birds, we reveal within-species variation in heterospecific social associations in follower species, which correlates with how individual followers are able to find novel food patches that are paired with social information from heterospecific vocalizations. Conversely, ‘nuclear’ species rate of finding novel food patches are correlated with number of connections to all flock members, conspecific and heterospecific. Our study provides evidence that individual variation in social connections within and across species influence foraging benefits in mixed-species groups.

## Introduction

Mixed-species groups are ubiquitous in nature across a broad taxonomic range and show remarkable variation in group size, species composition, species roles, seasonality, stability, and strength of associations (Goodale et al., 2017). Participation in these groups is generally attributed to two primary adaptive benefits: enhanced foraging efficiency and decreased predation risk (Morse, 1977). However, it is often challenging to assess actual changes to foraging efficiency or reduction in predation within wild mixed-species groups, and quantitative examples in the literature remain sparse (Goodale & Beauchamp, 2010). One reason for this is that the benefits of participation in mixed-species groups are highly context-dependent, shifting based on resource availability, predation risk, and the presence of particular group members. A central challenge to quantifying these benefits is that heterospecific information is not uniformly available to all members of a participating species; rather, its acquisition is dependent on individual interaction histories. Unraveling this complexity requires shifting analytical focus from the species-level to the individual-level (Farine et al., 2012; Vander Meiden et al., 2023). A focus on individual-level interactions enables us to evaluate the fitness consequences of social learning, link mixed-species behavior to trait evolution, and understand how this social topography scales up to broader ecological communities (Vander Meiden et al., 2023).

A defining characteristic of mixed-species assemblages is social information transfer, which can shape the local assembly and group movements as well as the spatial structure of the broader ecological community (Goodale & Beauchamp, 2010). There is a long line of evidence that adaptive benefits of mixed-species assemblages are linked to information transfer between species. For example, Krebs, (1973) showed that two species of small North American birds - black-capped chickadees (*Poecile atricapillus*) and chestnut-backed chickadees (*Poecile rufescens*) - mutually benefit from each other through interspecific social learning of foraging behavior. Within avian mixed-species flocks (hereafter MSFs), species roles are defined based on information production and use. According to this framework, nuclear species (leaders) typically function as information producers, whereas follower species exploit that information (Sridhar et al., 2009). For example, in Neotropical MSFs non-obligate ant-following species learn to eavesdrop on vocalizations of obligate ant-followers to locate army ant swarms (Pollock et al., 2017). Similarly, Carolina chickadees (*Poecile carolinensis*) seem to function as nuclear species in North American MSFs as their vocalizations attract individuals to foraging sites (Brooks & Freeberg, 2024; Freeberg & Lucas, 2002; Mahurin & Freeberg, 2009).

Previously, we examined the patterns of arrivals of North American MSFs of black-capped chickadees (*P. atricapillus*), white-breasted nuthatches (*Sitta carolinensis*), and downy woodpeckers (*Dryobates pubescens*) at novel feeders in relation to three playback treatments and found support for the role of black-capped chickadees in flock recruitment (Hafeez, Vander Meiden, Johnson, et al., 2026b). We found that chicka vocalizations of black-capped chickadees functioned as a strong conspecific recruitment signal, driving both faster and increased numbers of arrivals at novel feeders. While conspecific arrivals were clearly and strongly influenced by vocalizations of black-capped chickadees, the effect on arrivals of the follower species, white-breasted nuthatches and downy woodpeckers, was more nuanced. Follower species exhibited shorter latencies to arrival in response to chicka playbacks, but this effect was less pronounced. This weak effect can be attributed to asymmetry in benefits among species and variation among individuals of the follower species, i.e., if only a few nuthatches and woodpeckers gained the benefits of chickadee vocalizations (compared to many chickadees), then the overall effect at the species level would be weaker, even if some individuals gained large benefits. That is, a species-level analysis without accounting for individual variation could potentially cause the effects of heterospecific information to appear weaker than they really were.

Social networks can be used to quantify patterns of associations among individuals, and they have become a powerful tool for understanding patterns of information flow in animal societies (Duboscq et al., 2016; Tóth et al., 2020). However, although social networks are increasingly used to study such dynamics within MSFs (Bharadwaj et al., 2025; Farine et al., 2015; Richardson et al., 2022; Vander Meiden et al., 2023), mapping individual-level social interactions across multiple species presents logistical and conceptual challenges. For example, when individuals of multiple species associate, do heterospecific associations have quantitatively or qualitatively different effects on individuals as compared to conspecific associations? In the context of food patch discovery in European MSFs, previous research found an overall stronger effect of conspecific connections on information transmission as compared to heterospecific connections (Farine et al., 2015). This is one example of the potential use of social network analysis to test information theory in MSFs by measuring the foraging benefits of information gained through intraspecific and interspecific associations. Beyond single-layer analyses, emerging analytical frameworks like multilayer and multiplex networks offer promising avenues for disentangling complex species interactions, scaling seamlessly from individual associations to group-level information transmission (Finn et al., 2019; Hasenjager et al., 2021).

This study uses an individual-level social network framework to gain deeper insight into how social information provided by vocalizations of nuclear species (black-capped chickadees) and follower species (white-breasted nuthatches) influence food finding by MSF members. We hypothesized that novel feeder discovery is facilitated by individual-level social connectivity within these flocks. We hypothesized that, if foraging is socially facilitated, then individual birds that flock together at regular feeders will find novel feeders together. We further hypothesized that individuals that maintain more social connections will be more likely to find novel feeders. Finally, we hypothesized that, for nuthatches and woodpeckers, social connections to chickadees would influence their ability to find novel feeders paired with chicka vocalizations of black-capped chickadee, which was previously shown to attract chickadees to novel feeders (Hafeez, Vander Meiden, Johnson, et al., 2026b). Our findings reveal how fine-scale social associations shape the variable foraging advantages gained by individual flock members.

## Methods

### Study system

We conducted our study from 24 January 2021 to 25 February 2021 at Reller Prairie Field Station (40.6150^°^N, 96.7613^°^W) in Martell, Nebraska, USA. The site consists of a small deciduous woodland dominated by oak and other hardwood trees. Resident birds commonly form MSFs during the winter. These flocks typically include three core species: black-capped chickadees, white-breasted nuthatches, and downy woodpeckers. Other species may join these flocks occasionally, such as red-bellied woodpeckers (*Melanerpes carolinus*), brown creepers (*Certhia americana*), and red-breasted nuthatches (*Sitta canadensis*).

The three core MSF species were monitored at this site via RFID feeder system. Birds had access to eight stationary tube feeders (i.e., fixed feeder array) that were filled weekly with safflower seeds. Feeders were 54 m apart on average (i.e., the mean pairwise distance between closest fixed feeders). During fall, birds were opportunistically caught at these feeders using mist nets and fitted with USGS aluminum bands and passive integrated transponder (PIT) tags embedded in plastic leg bands (Eccel Technology Ltd., Groby, Leicester, UK). In the winter, the feeders were equipped with radio frequency identification (RFID) antennas, which continuously recorded daily feeding activity of MSFs. The RFID system operates through short-distance communication between reading units (antennas) and transponder (PIT) tags (Bridge et al., 2019). When PIT tags are activated by the antenna’s electromagnetic field, they transmit a tag-specific alphanumeric identification code which is recorded by the RFID data logger. We used an Arduino-based RFID data logger connected to a circular coil antenna (QKits Electronics, Ontario). Antennas were mounted on the landing perch in front of each feeder’s only opening and were programmed to scan for tags at regular intervals of every <u>^1^</u>th of a second throughout 24 hours. With each detection, the PIT tag ID, timestamp, and date were stored in the data logger. We used manual test tags at known times to verify clock accuracy across all RFID units and corrected for any discrepancies in time during data processing.

Feeders operated throughout the winter, continuously collecting data on feeder visitation, which was used to generate flocking associations (see below). In fall 2020, prior to the experiment described below, a total of 12 black-capped chickadees, 24 white-breasted nuthatches, and 20 downy woodpeckers had been banded with PIT tags at the site and were thus used to examine how individual social networks, generated from flocking behavior at the fixed feeders, impacted novel food finding.

### Field experiments

Our data come from novel feeder finding experiments that were previously analyzed at the species level to determine how vocalizations influenced recruitment of mixed-species flocking species to novel feeders (Hafeez, Vander Meiden, Johnson, et al., 2026b). We employed a matched-pair design consisting of 15 sets of two-hour long experiments in which three novel feeders were placed in random locations in the study site and each paired with a different acoustic stimulus (explained below). We conducted experiments during the hours between 08:00 and 13:00. To ensure unbiased placement, we determined novel feeder locations by randomly generating GPS coordinates within the study area. We maintained a mean pairwise distance of 197 m (64–404 m) between novel feeder sites. For each set of simultaneous trials, we randomly assigned playback treatments and exemplars to novel feeder locations. We monitored foraging recruitment at novel feeders during trials using the same RFID system described above.

To estimate PIT tag detection rates for each species, we cross-referenced video footage taken at novel feeders with automated data from RFID loggers on those same feeders (Hafeez, Vander Meiden, Johnson, et al., 2026b). The estimated PIT tag detection probabilities per visit were 0.58 for black-capped chickadees, 0.49 for downy woodpeckers, and 0.52 for white-breasted nuthatches.

### Playback stimuli

We used three different playback stimuli: chicka vocalizations of black-capped chickadee (as described in Ficken et al., 1978), quank calls of white-breasted nuthatch (as described in Ritchison, 1983), and white noise (as a control treatment). Our chicka stimuli consisted of individual A, B, or C notes from the chick-a-dee call complex in a randomized order with each note separated by natural inter-note intervals. Vocalizations used in playbacks were obtained from the xeno-canto database and recordings from a nearby location (Pioneers Park Nature Center [40.7761^°^N, 96.7691^°^W], Lincoln, NE; approximately 17 km from the study site). We constructed three unique exemplars for each stimulus to serve as replicates. For both species, we standardized playback rates at 10 vocalizations per minute. Playback stimuli were constructed by alternating one minute of treatment (bird vocalizations or white noise) with four minutes of silence. This five-minute sequence was repeated to produce two-hour audio files. We normalized bird vocalizations and adjusted playback amplitudes within the natural range of respective species (Nowicki, 1983; Podos & Patek, 2015; Rossetto et al., 2025). We produced white noise for the control stimulus in Raven Pro 1.6.5 (Cornell Lab of Ornithology). We used Axess SPBT1031 portable loudspeakers to broadcast playbacks at the novel feeder sites. To minimize the impact of human disturbance, a 10-minute silent interval was included at the start of each file. Further details of the playback stimuli are available in (Hafeez, Vander Meiden, Johnson, et al., 2026b).

### Statistical analysis

#### Testing whether the social network predicted which birds found novel feeders together

We used the RFID data stream from the fixed feeder array to construct the social network of associations (“fixed feeder network”). We restricted the data from the fixed feeders to dates spanning the first and last day of the food-finding experiments. In addition, we excluded 9 days during which many fixed feeders were non-operational due to extremely low temperatures, causing the lithium batteries to fail. This resulted in 16 days of data. From this dataset, we defined ‘flocks’ using the Gaussian Mixture Model (GMM) to detect ‘local gathering events’ or concurrent pulses of visits by tagged birds at a given feeder (Psorakis et al., 2012, 2015). We then calculated the association index between each pair of birds as the Simple Ratio Index (SRI) (Cairns & Schwager, 1987), which is the number of times individuals A and B were observed in the same flock, divided by the number of times A or B were seen in total. We implemented the Gaussian Mixture Model and calculated the association indices using the R package asnipe (Farine, 2013). We then used the igraph package to construct the social network (Csardi & Nepusz, 2006).

We used the RFID data stream from the experimental novel feeders to construct a ‘co-finding network’. We calculated flock associations at novel feeders using the same GMM method described above. In addition, we calculated the number of novel feeders discovered per playback treatment, as well as the cumulative number of novel feeders found by each PIT tagged bird throughout the study period.

To test the hypothesis that social associates facilitate foraging in MSFs, we compared the fixed feeder network and the co-finding network using a Multiple Regression Quadratic Assignment Procedure (MRQAP) with Double Semi-Partialing (DSP). This regression method provides robust estimates while accounting for the inherent non-independence in network data. We did this by testing whether the associations at fixed feeders predicted flock associations at novel feeders during the experimental trials.

#### Testing how social connections influence variation in individuals’ ability to find novel feeders

Next, we asked whether a bird’s connections in the social network predicted the number of novel feeders they found over the course of the experiment. We calculated degree centrality, or the number of connections that each bird had in the social network, as the measure of the individual’s social connectivity. We determined ‘degree to chickadee’, i.e., the number of connections to individual black-capped chickadees for each bird, to distinguish between the effects of general connections and the associations made with a nuclear, often informant species.

We also measured the variation in individuals’ rate of feeder use as a covariate in our models. We applied a 4-second threshold to distinguish separate ‘visits’ at the same feeder (Madsen et al., 2021). Any consecutive records of the same bird at the same feeder within this time window were considered as a single visit. We calculated the daily feeder visitation rate as the number of visits divided by the cumulative feeder-hours (the sum of the amount of time between first and last record at each feeder for a given day), and then calculated the mean daily visitation rate for each bird as a measure of their feeder use.

We used generalized linear models (GLMs) to test whether social network connections predict the number of novel feeders found by individuals. We first fit a global negative binomial GLM using the total number of novel feeders found as the response variable, and degree centrality, mean daily feeder visitation rate, species, and all interactions as fixed effects. Given that there was an interaction between the effects of degree centrality and species (*p* = 0.035), we then fit separate models for each species.

Next, we tested the hypothesis that social connections to black-capped chickadees influence the probability that members of the follower species would find novel feeders paired with chicka vocalizations. To test this hypothesis, we used the number of novel feeders found by each bird as the response variable, and the species, their degree to chickadees, and the mean daily visitation rate as fixed effects in negative binomial GLMs. Because the sample sizes were relatively low, we also conducted AIC-based model selection to determine whether a reduced model was a better fit to the data. We conducted this analysis for each playback treatment (chicka, quank, and control) and evaluated all models for overdispersion and zero-inflation. We visualized relationships using scatter plots and regression lines with 95% confidence intervals derived from model predictions.

## Results

At the fixed feeder array, we recorded 3,821 flocks of ≥2 birds (*n* = 5,245 flocks including individuals visiting alone) involving 48 tagged birds during the experimental period. From the novel feeder experiments, we recorded a total of 243 flocks of at least ≥2 birds (*n* = 408 flocks including individuals visiting alone) involving 34 unique individuals.

### (a) Flock associations at fixed feeders predicted co-arrival at novel feeders

The MRQAP results showed that the strength of flock associations at the fixed feeders strongly predicted flock associations at novel feeders (figure 1; estimate = 0.52, *F*_1,526_ = 341.0, *R*^2^ = 0.39, *p* < 0.001).

**Figure 1.**
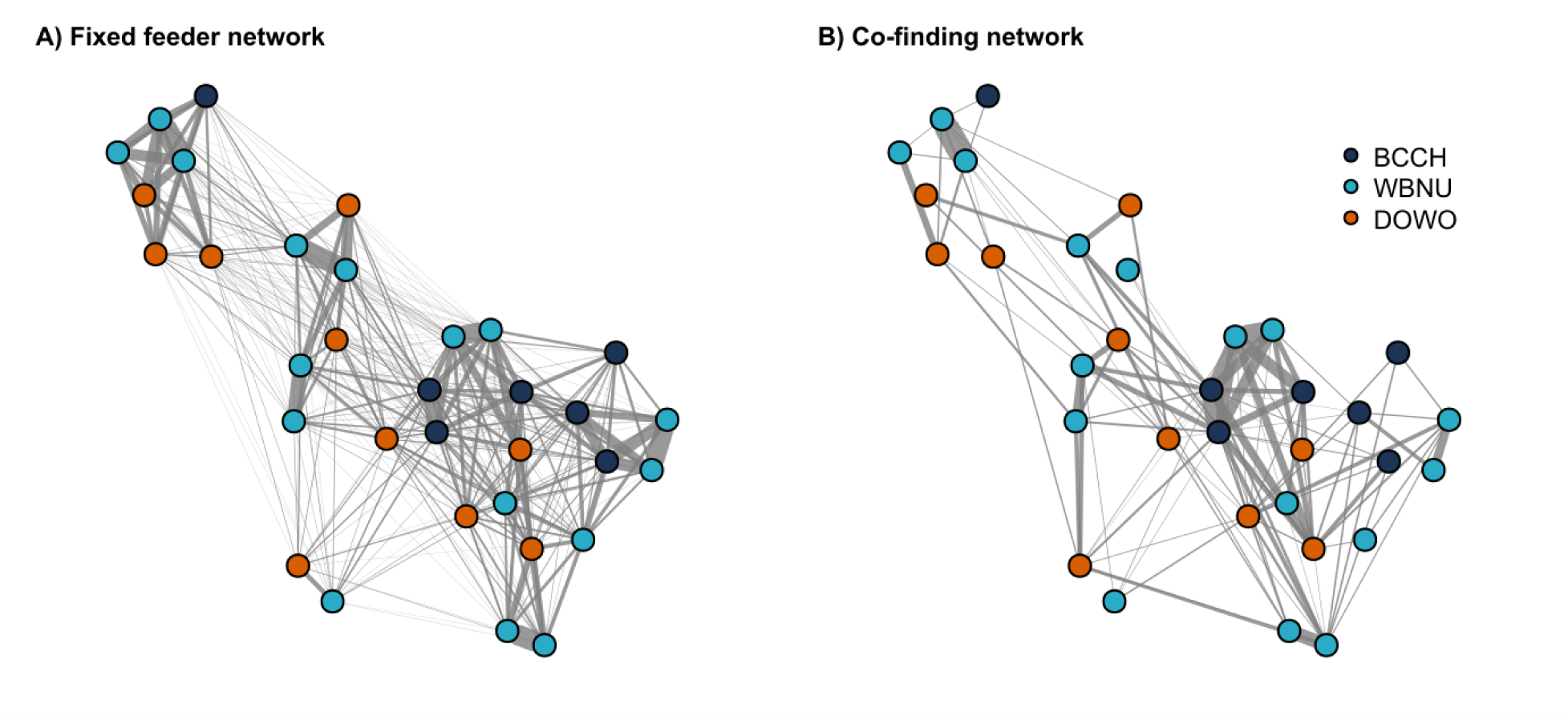
Social network structure of MSFs at fixed and novel feeders. Networks illustrate social associations constructed from (**A**) flock associations at fixed feeders and (**B**) flock associations at novel feeders during trials. Nodes represent individual birds, with each individual’s position fixed across both network panels to facilitate direct visual comparison. Nodes are coloured by species: black-capped chickadee (BCCH; P. atricapillus; dark blue), white-breasted nuthatch (WBNU; S. carolinensis; cyan), and downy woodpecker (DOWO; D. pubescens; orange). Edge width represents the strength of social associations. Pre-existing social associations at fixed feeders strongly predicted which individuals co-discovered novel feeders.

### (b) Chickadees that maintained more social connections were more likely to find novel feeders

The rate of discovery of novel feeders was not predicted by feeder visitation rate in any species (table 1). Separate species models revealed a strong positive effect of overall network degree centrality on the total number of feeders found by black-capped chickadees (*P. atricapillus*), but not by downy woodpeckers (*D. pubescens*) or white-breasted nuthatches (*S. carolinensis*) (table 1, figure 2).

**Figure 2.**
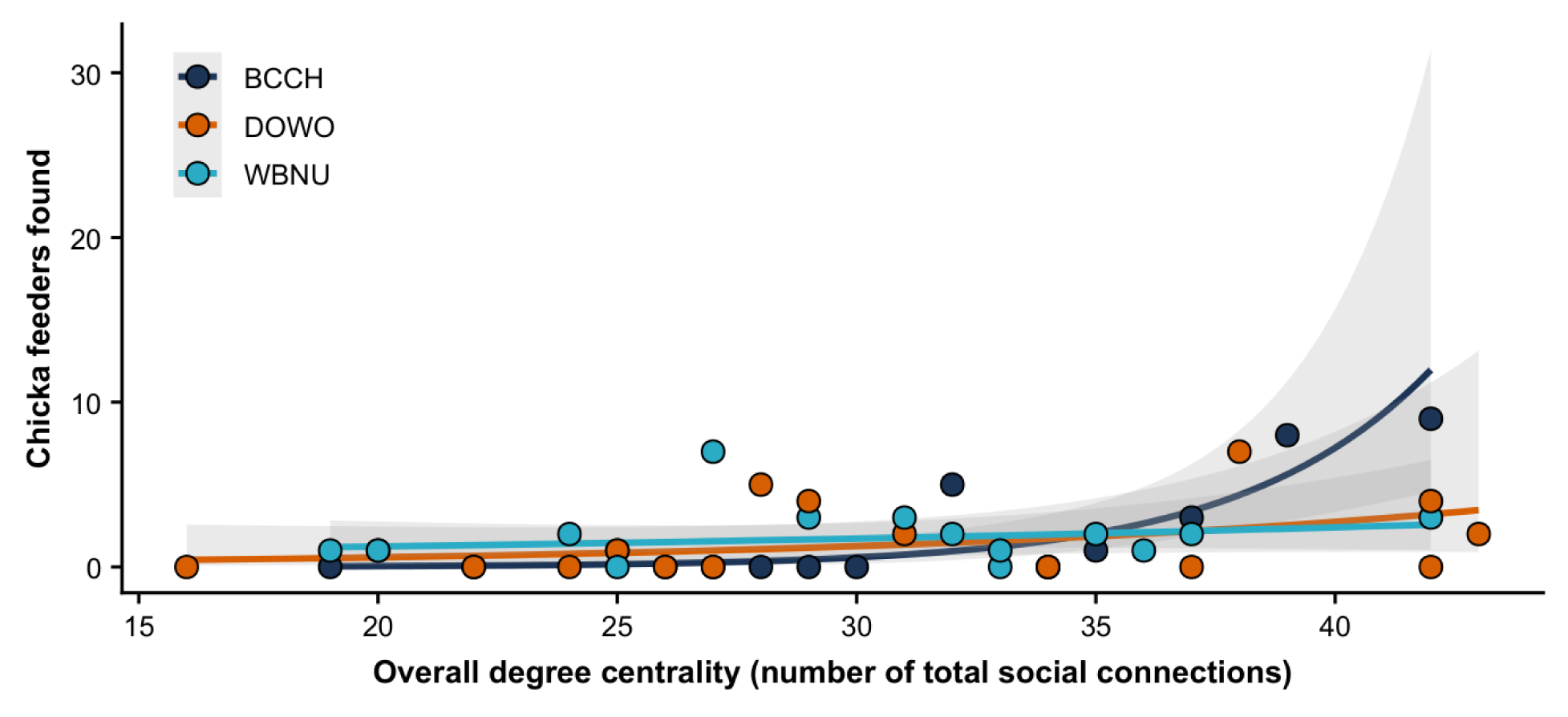
Relationship between overall social network centrality and novel feeder discovery across flocking species. Fit lines represent the predicted number of novel feeders found as a function of total degree centrality, with shaded grey bands indicating 95% confidence intervals. Points are colour-coded by species: black-capped chickadee (BCCH; P. atricapillus; dark blue), white-breasted nuthatch (WBNU; S. carolinensis; cyan), and downy woodpecker (DOWO; D. pubescens; orange). Total number of social connections significantly predicted foraging success only for the nuclear species, the black-capped chickadee.

**Table 1.** Species-specific negative binomial GLMs predicting the total number of novel feeders found by each species as a function of network centrality and mean daily feeder visitation rate.

| Species | Variable | Estimate ± s.e. | $z$ | $p$ |
| --- | --- | --- | --- | --- |
| **Black-capped chickadee** | **Degree centrality** | $0.18 \pm 0.06$ | 2.96 | **0.003** |
| (*P. atricapillus*) | Mean daily visit rate | $-0.79 \pm 1.64$ | -0.48 | 0.63 |
| Downy woodpecker | Degree centrality | $0.03 \pm 0.05$ | 0.61 | 0.55 |
| (*D. pubescens*) | Mean daily visit rate | $-0.06 \pm 0.93$ | -0.06 | 0.95 |
| White-breasted nuthatch | Degree centrality | $-0.006 \pm 0.03$ | -0.24 | 0.81 |
| (*S. carolinensis*) | Mean daily visit rate | $0.50 \pm 0.37$ | 1.34 | 0.18 |

### (c) Connections to chickadees improved food discovery for followers

While global degree centrality did not universally predict foraging success across all species (figure 2), we found that follower species’ benefits were contingent upon individually maintained social ties to nuclear species. Specifically, degree centrality to black-capped chickadees (*P. atricapillus*) had a very strong effect on the performance of both follower species combined (downy woodpecker and white-breasted nuthatch) in finding novel feeders when paired with the chicka playback treatment (figure 3). With each additional social connection to a nuclear flock mate, our model predicted a 23% increase in the number of chicka feeders found by both follower species (figure 3). The strongest AIC-supported model included only the number of connections to chickadees as a fixed effect. Conversely, we detected no effect of social connections to chickadees on the followers’ ability to find feeders paired with control or quank playback treatments (table 2).

**Figure 3.**
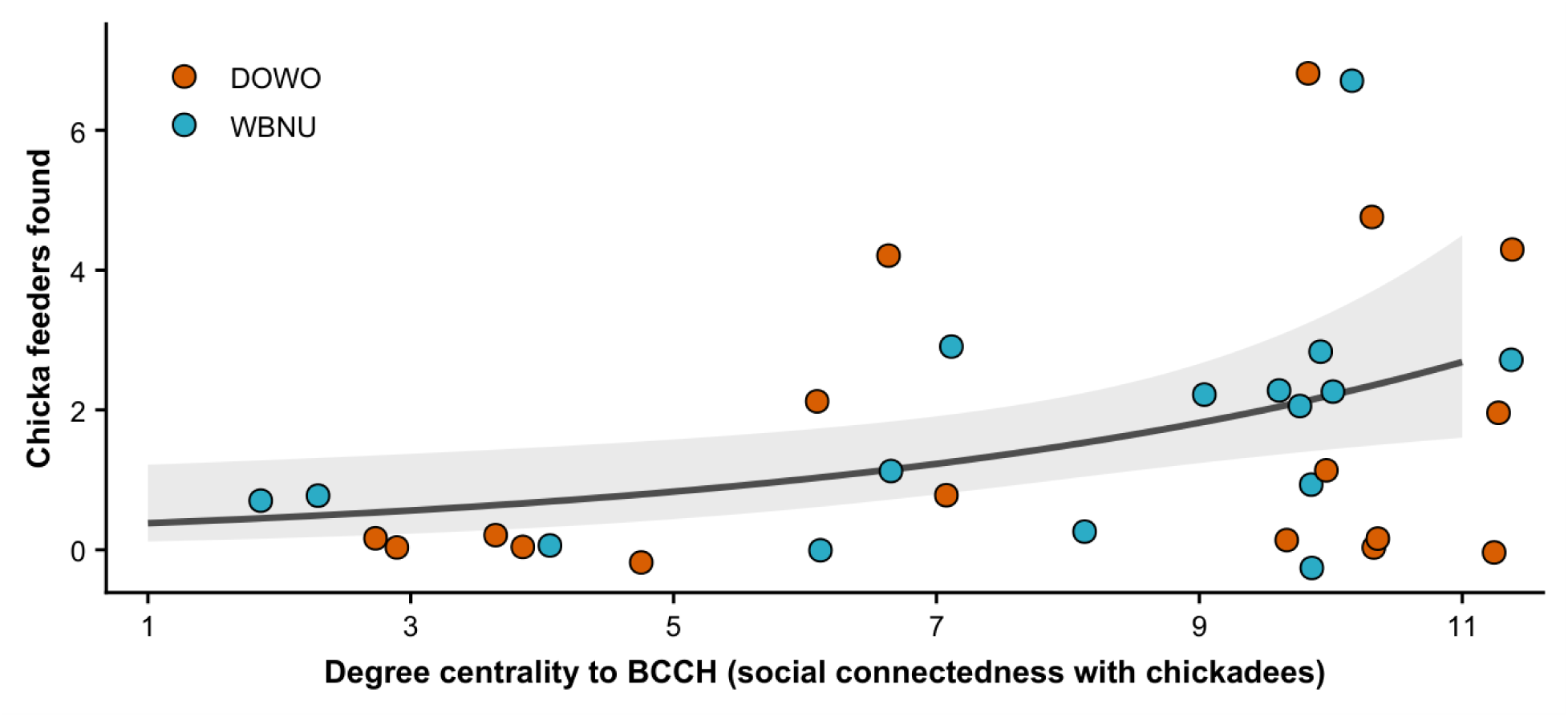
Heterospecific information use and foraging success of follower species relative to nuclear species associations. The fit line represents the predicted number of chicka feeders discovered by follower species as a function of their individual social connections (degree centrality) to black-capped chickadees (P. atricapillus). The shaded grey band indicates the 95% confidence intervals. Points denote individual follower birds, colour-coded by species: downy woodpecker (DOWO; D. pubescens; orange) and white-breasted nuthatch (WBNU; S. carolinensis; cyan). Direct social associations with nuclear species significantly increased feeder discovery rates exclusively when novel feeders were paired with relevant acoustic information (chicka playbacks).

**Table 2.** Negative binomial GLMs evaluating the effects of nuclear species associations and visitation rates on novel feeder discovery by follower species across playback treatments. Here, downy woodpecker is the reference group.

| Treatment | Variable | Estimate ± s.e. | $z$ | $p$ |
| --- | --- | --- | --- | --- |
| **Chicka playback** | **Degree to chickadee** | $0.19 \pm 0.08$ | 2.33 | **0.02** |
| | Species: white-breasted nuthatch | $0.26 \pm 0.45$ | 0.58 | 0.56 |
| | Mean daily visit rate | $0.17 \pm 0.49$ | 0.34 | 0.73 |
| Quank playback | Degree to chickadee | $-0.06 \pm 0.06$ | -1.02 | 0.31 |
| | Species: white-breasted nuthatch | $0.47 \pm 0.38$ | 1.23 | 0.22 |
| | Mean daily visit rate | $0.09 \pm 0.41$ | 0.23 | 0.82 |
| Control (white noise) | Degree to chickadee | $-0.06 \pm 0.06$ | -0.95 | 0.34 |
| | Species: white-breasted nuthatch | $0.53 \pm 0.38$ | 1.41 | 0.16 |
| | Mean daily visit rate | $0.60 \pm 0.41$ | 1.47 | 0.14 |

## Discussion

Our experimental results provide clear evidence to support the theoretical expectations that advantages of MSG formation can be context-dependent, asymmetric between species, and variable between individuals (Goodale et al., 2020; Vander Meiden et al., 2023). Our analysis revealed that the social network at regular feeders strongly predicted which individuals discovered novel feeders together as a flock, demonstrating that exploitation of new food resources by MSF members is facilitated by established social ties. Further, we found that the number of flock associates (both conspecific and heterospecific) at fixed feeders explained variation in novel feeder discovery rate among individuals in the nuclear species, black-capped chickadees. For white-breasted nuthatches and downy woodpeckers, the number of novel feeders discovered was predicted by the number of connections with black-capped chickadees, while connections to their own or the other follower species had no significant effect. Importantly, the effect of connectivity to chickadees was observed only at novel feeders paired with chicka playbacks, suggesting that social information transfer is a necessary mechanism for acquiring these benefits. Collectively, our results support the hypothesis that follower species in MSFs gain foraging benefits through direct connections with nuclear species, but that these benefits are not gained equally among members of follower species: individuals that regularly flock with the nuclear species (here, black-capped chickadees) are the ones that gain these benefits. Overall, our study highlights the importance of considering individual variation in the fitness consequences of species interactions.

Our results demonstrate that follower species can gain the foraging benefits of social information produced by chickadee vocalizations at novel feeders, but this advantage is primarily gained by individuals that regularly flock with chickadees. Thus, this individual-level analysis adds a new dimension to our previous study using this experiment, where we reported that chicka playbacks appeared to act as a strong recruitment signal primarily for conspecifics, while the effect on follower species appeared less pronounced (Hafeez, Vander Meiden, Johnson, et al., 2026b). Our current study reveals that this apparently weak effect among follower species was due to heterogeneity among individuals in their social connections to chickadees, a fine-scale pattern that did not emerge clearly when using species-level analyses. Because not all follower individuals maintained strong social ties with chickadees, nor were they uniformly attuned to utilizing heterospecific vocal cues, evaluating these dynamics at the species level obscured the individual-level benefits being achieved. Focusing on the individual-level social networks is therefore paramount for understanding the function of social information in MSFs; information transfer is fundamentally a transaction between individuals, and it is within this precise social framework that members of natural populations vary in their integrated sociality and accrued fitness benefits. Failing to account for this intricate variation not only masks crucial ecological patterns but constrains our broader understanding of how MSFs assemble, function, and evolve (Vander Meiden et al., 2023). Ultimately, our findings underscore that the adaptive benefits of MSF participation are not uniformly distributed at the species level, but are instead deeply rooted in unique social tendencies of individual flock members.

Our study contributes to a growing body of evidence demonstrating that fine-scale social relationships heavily influence foraging dynamics in mixed-species groups. Foundational research on winter-resident parid communities in the UK, comprising blue tit (*Cyanistes caeruleus*), great tit (*Parus major*) and marsh tit (*Poecile palustris*), established that the social network of flock membership at stationary feeders predicts arrival patterns at novel food patches (Aplin et al., 2015); and that both conspecific and heterospecific network ties contribute to the transmission of information about new feeding sites (Farine et al., 2015). We observed a similar pattern in our North American system: social associations at fixed feeders carried over to novel feeders, with strongly affiliated individuals arriving at novel foraging sites in close temporal proximity. This functional link between social association and collective exploitation of novel resources has now been documented across diverse taxonomic lineages, including group-foraging passerines (Aplin et al., 2012; Tóth et al., 2017), social primates (Hobaiter et al., 2014), and marine mammals (Allen et al., 2013). While these cross-taxa parallels underscore that stable social ties are highly influential, the precise foraging benefits derived from flocking can vary drastically based on a species’ functional role within the MSF as well as an individual’s degree of social connectivity.

In black-capped chickadees, we found a strong correlation between overall network centrality and novel feeder discovery rates. This link between social connectivity and individual foraging success could be explained by multiple, non-mutually exclusive mechanisms. It is possible that individuals with good food discovery skills are followed by a higher number of flock mates (Brown, 1986; Reebs, 1999), subsequently emerging as highly central nodes in the association network. This process is likely facilitated by vocal recruitment as proficient food finders may attract flock members to newly discovered resources - a behavioral strategy well-documented in chickadees and other parids (Mahurin & Freeberg, 2009; Suzuki, 2016). Alternatively, chickadees that flock with a higher number of individuals may gain advantages that increase their probability of discovering novel food. For instance, participation in larger groups can directly facilitate problem-solving (Cantor et al., 2020; Liker & Bókony, 2009; Morand-Ferron & Quinn, 2011), and maintaining diverse conspecific and heterospecific associations across flocks may grant individuals access to a larger cognitive toolkit and a richer pool of behavioral competence (Freeberg et al., 2017, 2024; Giraldeau & Caraco, 2000). This higher investment in interspecific connections may lead to more social information (Aplin et al., 2012; Farine et al., 2015). Taken together, our data demonstrates that maintaining high network centrality is an adaptive behavioral strategy that enhances foraging efficiency in black-capped chickadees, providing a buffer against environmental stressors during harsh winter conditions.

In white-breasted nuthatches and downy woodpeckers, we found a strong correlation between individual degree centrality to black-capped chickadees and novel feeder discovery rates under chicka playback conditions. This contrast between the influence of conspecific and heterospecific social ties highlights the intricate organization of individuals within MSFs. Black-capped chickadees are highly gregarious; they maintain cohesive conspecific social groups, produce complex vocal signals, and act as information producers or sentinels in MSFs, thus assuming the ‘nuclear species’ role (Sridhar et al., 2009). Indeed, parids seem to universally fulfill this nuclear function due to their structurally complex vocalizations that encode highly specific contextual information, such as local risk levels (Coppinger et al., 2020; Krams et al., 2012; Lucas & Freeberg, 2007). Meanwhile, follower species such as white-breasted nuthatches and downy woodpeckers scrounge on this public information, though our study demonstrates that the propensity to scrounge is not uniform, varying significantly among individuals within the population based on their proximity to nuclear species (Carlson et al., 2020; Sridhar et al., 2009; Sullivan, 1985). This individual tendency to scrounge on heterospecific information is likely tied to each follower’s social system. For the white-breasted nuthatch, a species with year-round territoriality, intense intraspecific aggression, and niche competition, interspecific sociality provides an evolutionary shortcut to flocking benefits without the costs of conspecific rivalry (Harrison & Whitehouse, 2011; Kilham, 1972). For downy woodpeckers, interspecific ties are important because their selective calling behavior makes conspecific social information less readily available (Sullivan, 1985). Our data suggest that individual nuthatches and woodpeckers that maintain proximity to chickadees can efficiently capitalize on information generated by them, offsetting personal foraging risks and driving flock formation (Farine et al., 2015; Sridhar et al., 2009). Viewed broadly, MSFs represent the aggregate interactions among individuals from diverse species, each governed by its own species-specific social system which dictates the functional roles that each species plays within the flock (Vander Meiden et al., 2023). These distinct ecological roles drive asymmetry in social information production and use, underscoring the dynamic, multi-layered architecture of MSFs as integrated social systems.

Our study reveals that, for follower species, social connections to conspecifics may serve qualitatively different functions than social connections to heterospecifics - specifically black-capped chickadees. Such qualitative differences in social connections within versus between species suggest complexity in the MSF social system that is difficult to capture in a typical social network. Such complexity can be captured by a multilayer network approach (Finn et al., 2019). Specifically, the individual-level, multi-species social network of MSFs can be represented as a ‘node-colored network’ (also known as ‘multitype network’; Kivela et al., 2014) in which the intraspecific network of each species is represented as a distinct layer, while interspecific connections are represented as intralayer edges (Finn et al., 2019). As an example, we reformat our fixed-feeder and co-finder networks as an interconnected multilayer network (figure 4), illustrating how connections within layers (conspecific) and across layers (heterospecific) vary among individuals in each species.

**Figure 4.**
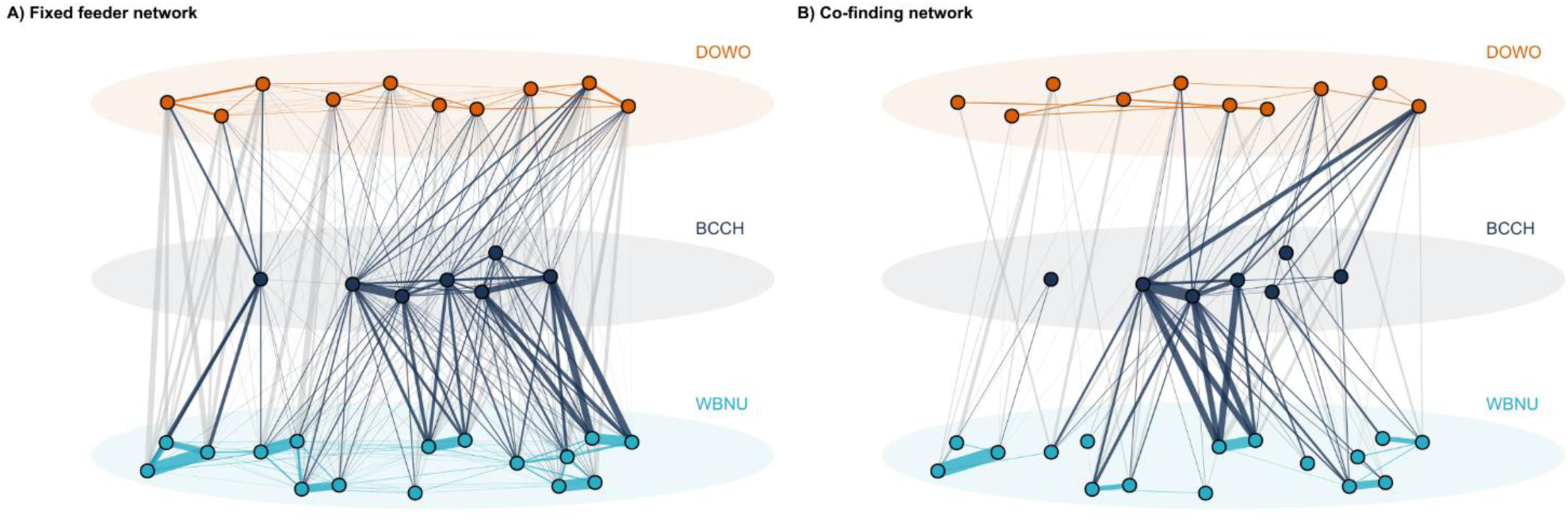
Multilayer network representation of social associations shown in figure 1. These interconnected multilayer networks illustrate MSF social associations constructed from (**A**) flock associations at fixed feeders and (**B**) flock associations at novel feeders during trials. Nodes represent individual birds, with each individual’s position fixed across both network panels to facilitate direct visual comparison. Nodes are stratified into parallel vertical layers by species: downy woodpecker (DOWO; D. pubescens; orange, top layer), black-capped chickadee (BCCH; P. atricapillus; dark blue, middle layer), and white-breasted nuthatch (WBNU; S. carolinensis; cyan, bottom layer). Edge width represents the strength of social associations within layers (intraspecific) and across layers (interspecific). Intraspecific edges (within species associations) match the corresponding species colors. Interspecific connections with nuclear species (BCCH-DOWO, BCCH-WBNU) are colored dark blue whereas interspecific connections between follower species (DOWO-WBNU) are rendered in light grey. The pattern of social associations exhibits remarkable variability within and across species. Also notice how within species associations change between fixed and co-finding networks.

This multilayer topology reveals structural asymmetry as a key evolutionary feature of these networks where nuclear species exert a disproportionately large influence on information flow across layers, yet the participation of follower species is critical for the collective foraging success of the flock. The within-species variation in heterospecific social associations correlated with individual followers’ ability to find novel feeders that were paired with social information from heterospecific vocalizations, which provides empirical evidence that individual variation in social connections within and across species influences foraging benefits gained in mixed-species groups. While analytical approaches of some types of multilayer networks, particularly multiplex networks (in which each layer represents a distinct interaction among the same sets of individuals) have begun to advance within animal social network studies (Finn et al., 2019), node-colored networks remain virtually unexplored. There is increasing evidence in many contexts that organisms gain benefits from associating with heterospecific individuals, but most studies fail to account for individual-level variation in the costs and benefits of such species interactions. Our study, together with (Farine et al., 2015), demonstrates that *a)* mixed-species groups may be an ideal system for advancing this branch of network analysis and *b)* quantifying fitness consequences of mixed-species social associations at the individual-level is important for understanding complex social relationships and mechanisms underlying community assemblages. Finally, our study highlights that individuals can navigate an intricate, interspecific social landscape to survive environmental challenges.

In conclusion, our findings empirically quantify the fine-scale asymmetries that underpin avian MSFs. While our findings stem from a single study population tracked over a single winter season, we provide high-resolution empirical evidence of socially mediated foraging benefits within MSFs. Moving forward, it will be very insightful to gather longitudinal data to evaluate how long-term demographic factors (such as age, sex, and individual turnover) alongside inter-annual climatic variations modulate the seasonal stability of these multilayered social structures. Expanding this individual-level analytical framework to diverse ecological contexts will be crucial for determining the ubiquity of complex multi-species networks that underlie much of the species interactions affecting ecological communities.

## Ethics

All experiments were conducted with approvals from all appropriate governing bodies regarding research ethics: the University of Nebraska-Lincoln Institutional Animal Care and Use Committee (protocol no. 2062), U.S. Geological Survey (Federal bird banding permit no. 23759), and the Nebraska Game and Parks (permit no. 1196). We adhered to the guidelines for the use of wild birds in research set forth by the Ornithological Council (Fair et al., 2023).

## Data Accessibility

The compiled dataset and supplementary materials are publicly available via Harvard Dataverse online (Hafeez, Vander Meiden, Hand, et al., 2026). Baseline behavioral metrics and the compressed acoustic playback stimuli were published alongside (Hafeez, Vander Meiden, Johnson, et al., 2026b) and are available online via OSF (Hafeez, Vander Meiden, Johnson, et al., 2026a).

## Declaration of AI use

We used the large language model Gemini (Google) to assist with R code optimization for multilayer social network figures. The authors reviewed and verified all AI-generated output and take full intellectual and ethical responsibility for the content, accuracy, and integrity of the final manuscript.

## Authors’ Contributions

F.H.: conceptualization, methodology, investigation, data curation, formal analysis, visualization, writing-original draft, writing-review and editing; L.N.V.M.: conceptualization, methodology, investigation, writing-review and editing; A.E.H.: formal analysis, visualization, writing-review and editing; A.E.J.: conceptualization, methodology, investigation, supervision, writing-review and editing; S.F.U.: investigation, writing-review and editing; S.R.S.: data curation; D.S.: conceptualization, methodology, formal analysis, visualization, resources, supervision, funding acquisition, writing-review and editing. All authors gave final approval for publication and agree to be held accountable for the work therein.

## Supplements

Supplementary materials are available online at: https://doi.org/10.7910/DVN/914YP6

## Acknowledgements

We would like to thank the School of Biological Sciences and the School of Global Integrative Studies at the University of Nebraska-Lincoln for granting us access to the Reller Prairie Field Station. We would also like to thank Furqan Mahdi, Brady Pham, and Yasmin Worth, who contributed to the transcription of videotapes, as well as all lab affiliates who helped with bird banding. We also acknowledge Ehtisham-ul-Haq for his technical and moral support to F.H.

